# Improving pairwise comparison of protein sequences with domain co-occurrence

**DOI:** 10.1101/115543

**Authors:** Christophe Menichelli, Olivier Gascuel, Laurent Bréhélin

## Abstract

**Motivation:** Comparing and aligning protein sequences is an essential task in bioinformatics. More specifically, local alignment tools like BLAST are widely used for identifying conserved protein sub-sequences, which likely correspond to protein domains or functional motifs. However, to limit the number of false positives, these tools are used with stringent sequence-similarity thresholds and hence can miss several hits, especially for species that are phylogenetically distant from reference organisms. A solution to this problem is then to integrate additional contextual information to the procedure.

**Results:** Here, we propose to use domain co-occurrence to increase the sensitivity of pairwise sequence comparisons. Domain co-occurrence is a strong feature of proteins, since most protein domains tend to appear with a limited number of other domains on the same protein. We propose a method to take this information into account in a typical BLAST analysis and to construct new domain families on the basis of these results. We used *Plasmodium falciparum* as a case study to evaluate our method. The experimental findings showed an increase of 16% of the number of significant BLAST hits and an increase of 28% of the proteome area that can be covered with a domain. Our method identified 2473 new domains for which, in most cases, no model of the Pfam database could be linked. Moreover, our study of the quality of the new domains in terms of alignment and physicochemical properties show that they are close to that of standard Pfam domains.

**Availability:** Software implementing the proposed approach and the Supplementary Data are available at: https://gite.lirmm.fr/menichelli/pairwise-comparison-with-cooccurrence

## Background

Proteins are macromolecules essential for the structuring and functioning of living cells. Proteins generally have different functional regions which are conserved along evolution (Zmasek and Godzik, 2011) and are commonly termed as “functional motifs” or “domains”. Domains/motifs are found in different proteins and combinations (Bornberg-Bauer and Albà, 2013) and, as such, are functional protein subunits above the raw amino-acid level. Domain identification is thus an essential task in bioinformatics.

Two kinds of approaches can be used to identify these regions of a target protein. Profile analysis, also known as sequence-profile comparison, is a powerful method. This *non ab initio* method requires a database of protein domains. Pfam is one of the most widely used databases (Finn et al, 2016). In this database, each family of domains is defined from a manually selected and aligned set of protein sequences, which is used to learn a profile hidden Markov model (HMM) of the domain. To identify protein domains, each HMM of the database is used to compute a score that measures the similarity between the sequence and the domain. If the score is above a predefined threshold, the presence of the domain in the protein can be asserted. However this method may miss several domains when applied to an organism that is phylogenetically distant from the species used to train the HMM. This may happen for two reasons. First, if the protein sequence has encountered many evolution events, the HMM of the database may poorly fit the sequence specificity of the distant organism. Different approaches can be used in this case; see for example (Terrapon et al, 2012) for a few solutions to this problem. Another possibility is that some domains of the query organism are simply absent from the database. Databases like Pfam were built with eukaryotic sequences that originate mostly from plants, fungi and animals, and very few from the other groups. Hence, the proportion of proteins covered by a Pfam domain in plants, fungi and animals is around twice that found in the other super-groups on the eukaryote tree (Chromalveolates and Excavates) (Ghouila et al, 2014). For example, in *Plasmodium falciparum,* which is the organism responsible for the deadliest form of malaria, only 22% of its protein residues are covered by a Pfam domain while this percentage is as high as 44% for both yeast (*Saccharomyces cerevisiae*) and humans.

An alternative approach for identifying protein domains is to run an *ab initio* approach based on sequence-sequence comparison using pairwise comparison tools like FASTA (Pearson and Lipman, 1988) or BLAST (Altschul et al, 1990). These tools look for local similarities between a query protein and a sequence database like Uniprot (The UniProt Consortium, 2015). Because domains are sub-sequences conserved throughout evolution, local similarities between proteins usually correspond to these regions. Tools like BLAST use specific scoring functions for assessing similarities, and provide estimates of p-values (and e-values) under specific score distribution hypotheses. As sequence-sequence approaches do not include information from other homologous sequences, they may be more prone to false positives than sequence-profile approaches. Therefore, they are usually used with stringent score thresholds and hence may also miss several homologies. Different versions of BLAST were developed to improve the sensitivity. For example, PSI-BLAST (Altschul, 1997), which constructs a position-specific score matrix (PSSM) to perform incremental searches, PHI-BLAST (Zhang et al, 1998), which uses a motif to initiate hits, or DELTA-BLAST (Boratyn et al, 2012), which searches a database of pre-constructed PSSMs before searching a protein-sequence database to yield better homology detection.

Surprisingly, so far domain co-occurrence has not been used to improve the sensitivity of sequence-sequence approaches in proteins. Domain co-occurrence is a strong feature of proteins, based on the fact that many protein domains tend to appear with a limited number of other domains on the same protein (Bornberg-Bauer and Albà, 2013). A well known example are domains PAZ and PIWI, which are frequently found together (see Figure 1): when assessing proteins with the PAZ domain, the PIWI domain is frequently found. Functional studies have shown that domains that co-occur in proteins are more likely to display similar functions than domains that appear in separate proteins (Ye, 2004). Co-occurrence information has already been used for improving the sensitivity of sequence-profile approaches (Terrapon et al, 2009). However, it could also be of great help for sequence-sequence homology detection. For example, Figure 2 reports homologies found between a *Plasmodium falciparum* protein and three proteins from Uniprot. Most of these hits have moderate e-values and, taken independently, cannot be considered with high confidence. However, every hit actually co-occurs with one or two other hits on the same protein, and these co-occurrences are present in all three proteins. Taken altogether, this information adds strong evidence on the identified homologies.

**Figure 1.**
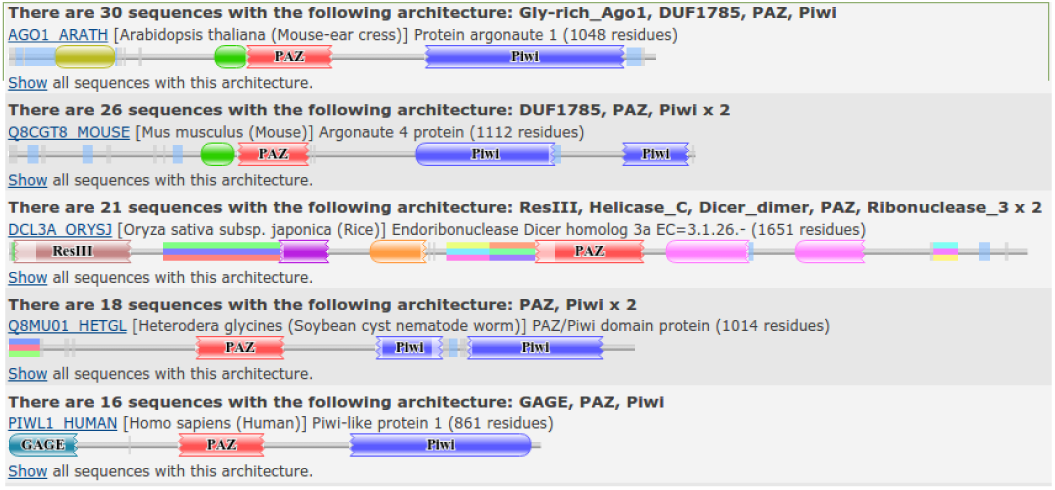
The five most common domain architectures involving the PAZ domain.

**Figure 2.**
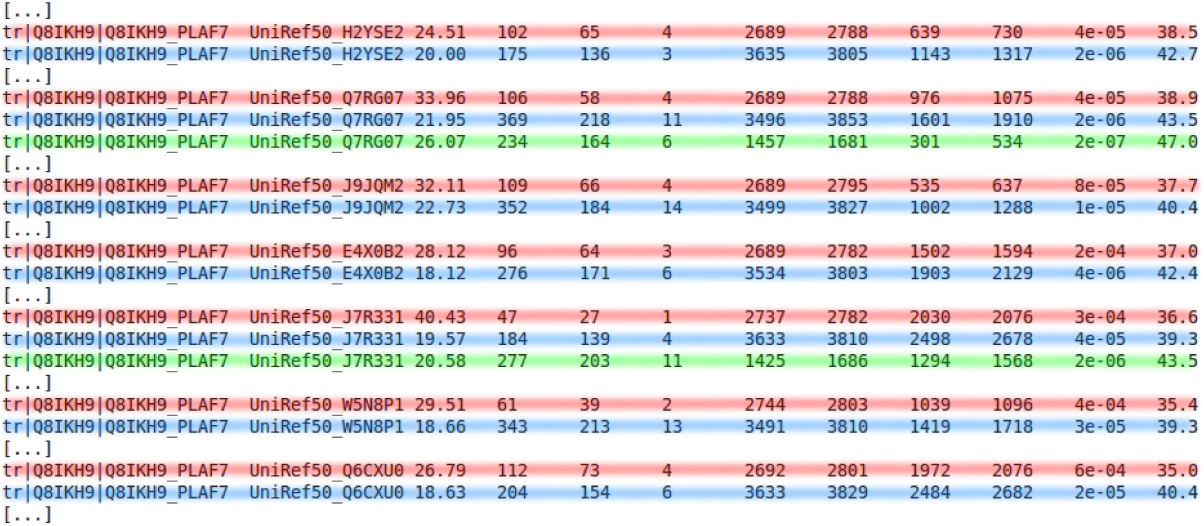
Extract of BLAST results on query sequence Q8IKH9_PLAF7 on UniRef50 (fields: query id, subject id, % identity, alignment length, mismatches, gap opens, q. start, q. end, s. start, s. end, e-value, bit score). Note that some hits are hidden for clarity. Depending on the target protein, the BLAST result reveals the co-occurrence of two/three independent sub-sequences (each sub-sequence is highlighted with a different color).

Here we propose a new method to take co-occurrence into account in a typical BLAST analysis and to construct new domain families based on these results. Our procedure is based on the analysis of co-occurring hit density along the query protein. We designed a procedure that uses this density to identify hit clusters that sign domain boundaries. We present our approach and propose a statistical test to assess the relevance of the identified clusters. Finally, we apply our approach to the entire *P. falciparum* proteome and show that, on this organism, it allows us to increase the number of significant hits by 16%. Moreover, our procedure for identifying new domain families enables us to increase proteome area that can be covered with a domain by 28%.

## Results

Our aim is to improve the sensitivity of pairwise comparison tools such as BLAST using co-occurrence information. The core of our approach is a new scoring function that takes co-occurrence information into account for assessing BLAST hits. This new scoring function allows us to identify interesting hits that would not be considered solely on the basis of BLAST results because of too high e-values.

### Discovering domains from BLAST results

The initial step is to perform a BLAST search of the query protein against a protein sequence database. In the experiments below, we used the UniRef50 database and the BLASTP software package (Altschul et al, 1990) with default parameters and a max e-value set at 10^−2^. This search gives us a list of all similarities found for the query sequence. Each pair of similar sub-sequences is called a hit. All hits smaller than 30 residues are removed, and in case of overlapping hits on a target protein only the hit with the lowest e-value is considered. Note that BLAST controls sequence complexity by automatically discarding low-complexity sub-sequences (default parameters).

Next we use the BLAST results to discover potential domains. Our hypothesis is that domains are the most conserved protein sub-sequences. Under this hypothesis, domains should correspond to regions with the highest number of hits in a classical BLAST search. Hence, we use the density of hits per residue to identify the potential domains of a given protein. The main drawback of this approach is that it can potentially produce many false positives. First, high peaks can also be due to low complexity regions that have not been properly masked by BLAST. Second, even if a peak corresponds to a domain, all hits that compose the peak are not necessarily true occurrences of this domain, and the proportion of contaminants can be high in certain cases. To avoid a maximum of false positives, our solution is to look at hits for which we have found a co-occurring hit. We define a co-occurring hit as another hit involving the same query and target proteins and that does not overlap with the former hit on any two proteins. Hence, rather than working on the hit density, we work on the co-occurring hit density, i.e. the density of hits with a co-occurring hit (see Figure 3). The goal is then to identify clusters of homologous hits that would represent protein domains. This is a difficult problem because all hits in a peak do not have the same length and are not perfectly aligned. Moreover, two adjacent domains on the query protein can also be adjacent in certain target proteins. In this case, the two domains would appear as a single long hit and may create ambiguity.

**Figure 3.**
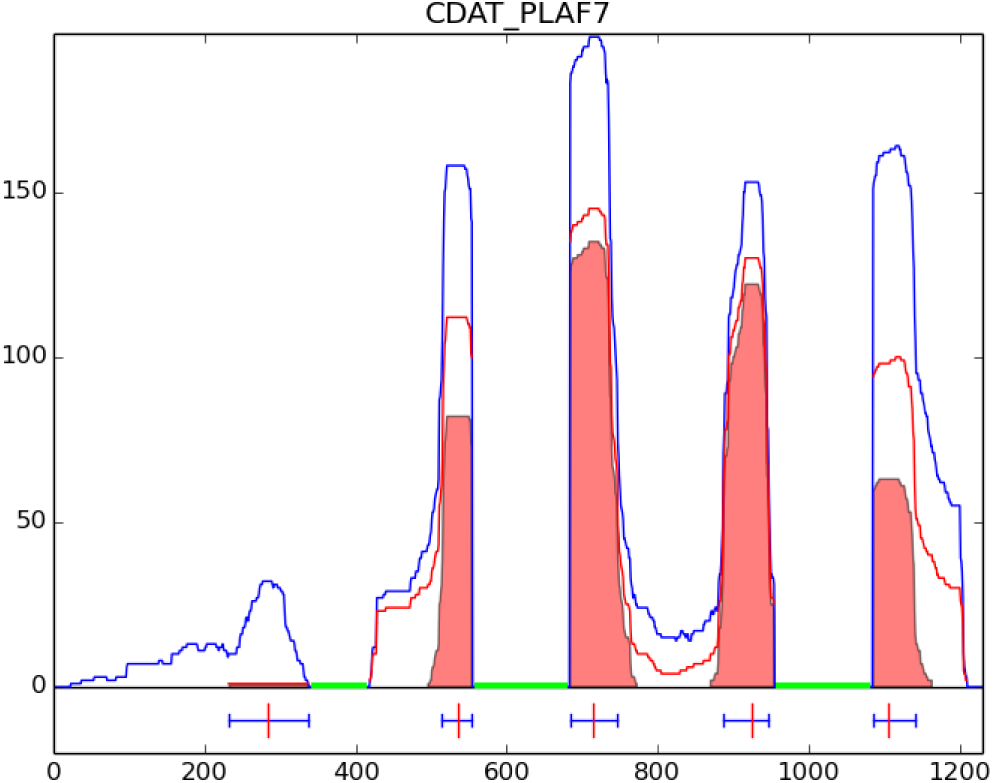
Density of BLAST hits per residue on the sequence CDAT_PLAF7. The blue line represents the density obtained using all hits. The red line represents the density obtained using only hits that have a co-occurring hit on the same protein. The filled regions in red show hit clusters identified by our method. Lines under the horizontal axis indicate positions of clusters on this protein. In this example, regions already covered by a Pfam domain were masked (green regions) so that no hits were identified by BLAST.

We developed an iterative heuristic for this problem. The method focuses on the co-occurring hit density, and starts by identifying the position associated with the highest peak. All hits covering this specific residue are selected and used to define the boundaries of the domain. This is done by iteratively selecting an homogeneous subset of these hits, *i.e.* by incrementally removing hits whose begin and end positions are too far from the other selected hits (see Methods for details on the algorithm). At the end of this process we identify a cluster of similar hits, *i.e.* similar in context because they share the co-occurrence property, and similar in sequence because they are located on the same region on the query protein. The cluster defines a protein domain family, with the different hits being different occurrences of the domain in different proteins. The region covered by this domain is then masked, and the whole procedure is resumed until no new domain can be identified on the protein.

### Evaluating cluster relevance

Selecting only hits that have a co-occurring hit should allow us to avoid many false-positives. However, co-occurrence can also be detected simply by chance. To control this, for each cluster (domain family), we compare the number of hits with co-occurring hits to the total number of hits in the cluster, and we estimate the probability of observing as many co-occurring hits in a cluster by chance. We use a binomial test for this purpose, and the p-value is computed as:

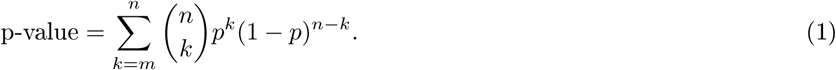

*m* is the number of target proteins with a hit in the cluster and a co-occurring hit outside the cluster; it is given by the red curve of Figure 3. *n* is the total number of target proteins with a hit in the cluster; it is given by the blue curve of Figure 3. *p* is the prior probability that a hit in the cluster has a co-occurring hit outside the cluster, given the total number of hits outside the cluster. This probability depends on the query protein and on the cluster. For example, some proteins may-have several low complexity regions with a lot of hits on UniRef50. In this case, it is easier for a hit in a given cluster to have co-occurring hits outside the cluster. Prior probability *p* is estimated by the total number of proteins with a hit on the query protein outside the cluster, divided by the total number of proteins in the database. It is conveniently computed by 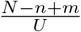, with *N* being the total number of proteins with a hit on the query protein and *U* the total number of proteins in the database (UniRef50). The number of proteins overlapping a cluster are computed on the position with the highest density of hits. If several residues have the same density, the central position is selected. In the following, clusters with a p-value higher than 10^−2^ are discarded.

### Estimating the number of false discoveries

The procedure described above is intended to be applied to all proteins of a given organism. For each discovered domain, the computed p-values allows us to check that the number of co-occurring hits cannot be found by mere chance. This ensures that the hits that compose a cluster likely occur from the same domain family. However, this does not ensure that the query protein is homologous to this particular family. Indeed, a protein may fortuitously possess two regions that resemble two domains that are frequently found together. In this case, we may observe many co-occurring hits, but the protein regions will not be homologous to the identified domain families. Although this should rarely happen, it is important to estimate the number of false positives detected by our procedure when it is applied to a whole proteome.

We designed a statistical procedure for this purpose. First, our approach is run on all proteins of the query organism, and the number of domains below a given p-value threshold is computed. Then all BLAST hits are randomly shuffled among all proteins, and the procedure is resumed on these random data. By comparing the number of domains “identified” in the random data to the number of domains identified in real data, we can get an estimate of the proportion of false positives of our procedure. However, it is important to preserve the homology relationship between hits during the randomization process. Hence, the hits are not shuffled independently among the proteins but by entire clusters (see Methods). Once the hits have been randomly distributed on proteins, the number of co-occurring hits are computed in each cluster, and a p-value is estimated as above. The number of clusters below the chosen p-value threshold is computed, and the entire procedure is resumed several times (*e.g.* 10 times) to get a better estimate of the number of domains that can be identified in random data. This number is then used to estimate the FDR of the procedure:

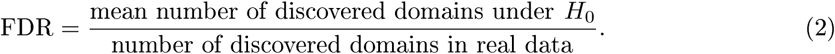

### Learning new models

Once new domain families have been identified with the above procedure, an additional and optional step is to learn an HMM for each cluster. Only clusters with at least 5 sequences are considered to learn a model. We first realigned the sequences of each cluster using MUSCLE software (Edgar, 2004), with default parameters. Flanking positions with less than 75% residues were removed. Then, each multiple sequence alignment (MSA) was used to train a HMM representing the corresponding domain family using HMMER (Eddy, 1998) (see Methods).

### Integrating known domains into the procedure

In order to focus on regions that are not already covered by a known domain, an interesting improvement is to include known domain information in the procedure. This is done at two levels. First, prior to the BLAST search, regions already covered by a Pfam domain are masked by replacing the covered residues by the unknown residue (X) which is ignored by BLAST. Next, when searching for co-occurring hits, known Pfam domains are also considered as potential co-occurring hits. Namely, if a query protein A has a BLAST hit on a target protein B, and if both A and B have the same Pfam domain D (and that D does not overlap the considered hit on A and B), then the BLAST hit is considered as having a co-occurring hit. Integrating known domain information has two advantages. First, it enables us to speed up the BLAST search by masking part of the query sequences. Second, by integrating accurate homology information, it also minimizes the chance of detecting false positive co-occurrences and improves the quality of the results.

### Experiments on *P. falciparum* proteins

We applied our procedure to *Plasmodium falciparum* proteins. Each protein sequence (release February 21 2016, on Uniprot) was used to run a BLAST search against the UniRef50 database (October 2015) (Suzek et al, 2015) restricted to eukaryotic sequences, which gathers 2 784 993 sequences from 2753 reference organisms with a maximum of 50% similarity between each pair of sequences.

We first ran our approach without integrating the known Pfam domains in the procedure (see Table 1). We identified a total of 19 100 clusters (4 633 with at least 5 hits) distributed on 4 394 different proteins (1915 with a cluster with at least 5 hits). Among the 19 100 clusters, 9 337 have a p-value below 1% (3 531 with at least 5 hits) on 2 633 different proteins (1 524 with a cluster with at least 5 hits). These clusters cover 37.96% of residues in the *P. falciparum* proteome. Clusters with at least 5 hits cover 13.87% of residues. The FDR of the procedure is estimated at 6%. Among the 3 531 clusters with at least 5 hits, 2 286 overlap an already known Pfam domain—only strong overlaps, *i.e.* greater than one third of the smallest domain, are considered here. We then assessed the extent to which our automatic procedure is able to recover the domains of the Pfam database. To answer this question, we generated the HMM associated with the 3 531 clusters and we compared this HMM to that of the overlapping Pfam domain. We used HHPred software (Soding et al, 2005) for this purpose (see Methods). Given two HMMs, this tool computes a local alignment of the HMMs with an associated p-value. From the HMM alignment, we also computed an overlap ratio by taking the ratio between the overlap length and the size of the longest HMM into account (see Figure 4). In most cases (87%), the HMM/HMM alignment had a significant p-value below 10^-10^, indicating that our HMM resembles (at least locally) the Pfam HMM. Moreover in 54% of cases, the HMM obtained with our cluster has an overlap with the Pfam HMM greater than 80%.

**Figure 4.**
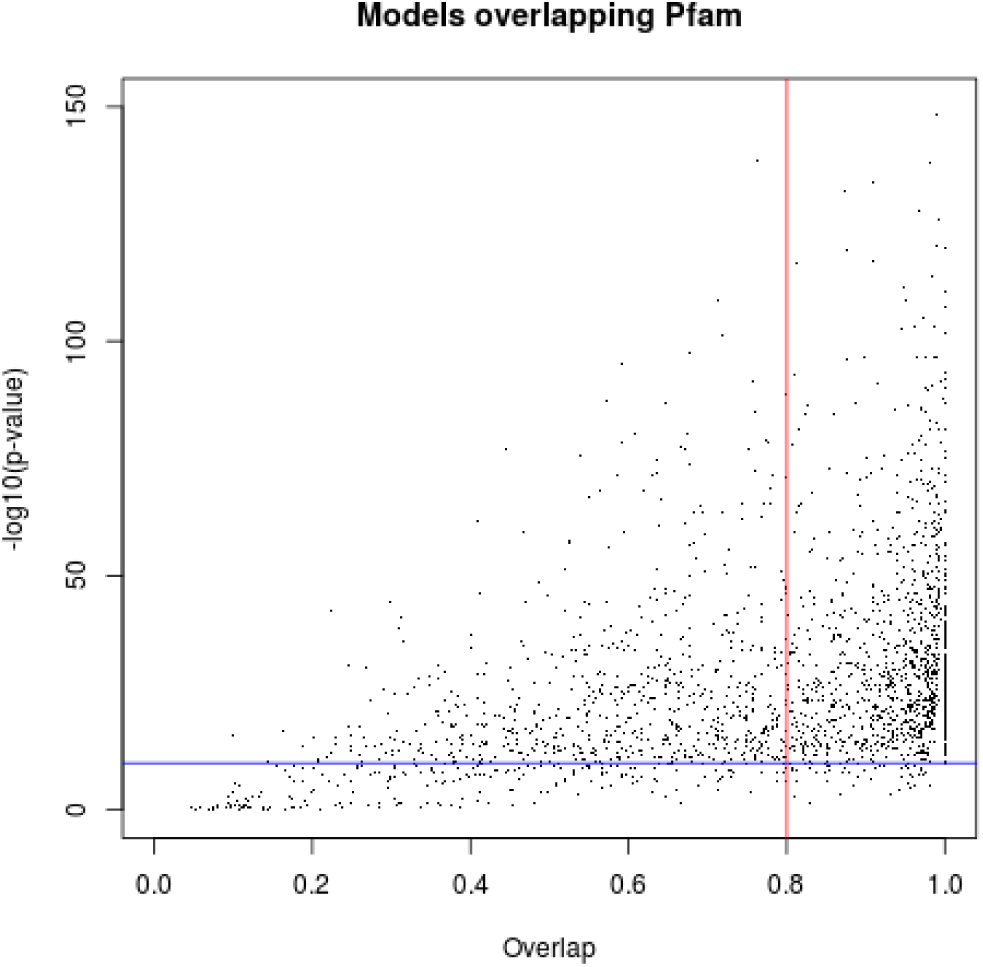
HMM/HMM comparison of domains identified by our approach that overlap a known Pfam domain. The x-axis shows the overlap ratio of the local alignment; the y-axis indicates the negative log of the alignment p-value; the blue line denotes the 10^−10^ p-value, while the red line denotes the 80% overlap.

**Table 1.** Summary of the number of new domain occurrences identified by the approach. In the “Without Pfam integration” experiment, analyses were done on the entire genome, without masking the already known Pfam domain occurrences in the BLAST search nor using them as potential co-occuring hits in the co-occurrence analysis. On the contrary, in the “With Pfam integration” experiment the already known Pfam domain occurrences were masked in the BLAST search and used as potential co-occuring hits in the co-occurrence analysis.

| Experiments | Without Pfam integration |  | With Pfam integration |  |
| --- | --- | --- | --- | --- |
| | all clusters | clusters $\geq 5$ hits | all clusters | clusters $\geq 5$ hits |
| number of clusters | 19 100 | 4 633 | 10 442 | 2 572 |
| with p-value $\leq 1\%$ | 9 337 | 3 531 | 8 530 | 2 473 |
| proteins involved | 4 394 | 1 915 | 3 039 | 1 378 |
| residue coverage | 37.96% | 13.87% | 28.28% | 7.12% |
| estimated FDR | 6% |  | 5.15% |  |

We next challenged our approach for identifying new domains not already covered by a Pfam domain. As explained above, we included all known Pfam domains in the procedure, *i.e.* all *P. falciparum* protein regions covered by a Pfam domain were masked before the BLAST search and were used as potential co-occurring hits during the co-occurrence analysis. This new analysis allowed us to identify a total of 10442 clusters (2 572 with at least 5 hits) distributed on 3 680 different proteins (1 430 with a cluster with at least 5 hits - see Table 1). Among the 10442 clusters, 8530 have a p-value below 1% (2 473 with at least 5 hits) on 3 039 different proteins (1 378 with a cluster with at least 5 hits). The MSAs of these domain families, and the HMMs that can be learned from the 2473 families with at least 5 hits are provided in the Supplementary Data. These clusters (which are not covered by Pfam domains) cover 28.28% of the residues in the *P. falciparum* proteome. Clusters with at least 5 hits cover 7.12% of residues. For comparison, Pfam domains cover 22% of this proteome. The FDR of the procedure is estimated at 5.15%. Clusters with a p-value < 1% gather a total of 335 311 hits. Among these, 46 960 have moderate e-values (> 10^−6^) and might not be considered in a classical whole-genome BLAST analysis. For comparison, the total number of BLAST hits with an e-value below 10^−6^ is 288 351. Hence, with the 10^−6^ threshold, the co-occurrence property allowed us to increase the number of considered hits by 16%.

We first investigated the origin of the hits selected by our procedure, compared to all BLAST hits and the whole Uniref50 database. Each hit was classified according to its target species in the five super-groups covering Eukaryota: Chromalveolata, Excavata, Rhizaria, Unikont, and Viridiplantae (Keeling et al, 2005). Figure 5.(a) reports the species distributions in all BLAST hits, only in the hits selected by our procedure, and in all proteins of the Uniref50 database. We observed slight enrichment of Chromalveolates in the BLAST hits compared to the Uniref50 database, and substantial enrichment of this super-group in the hits selected by co-occurrence. This was somewhat expected, considering that the proportion of false positives is likely lower for hits in the same super-group as *P. falciparum* than for hits outside this super-group. Moreover, this may also indicate that some of the newly discovered domains are specific to Chromalveolates. To assess this hypothesis, we computed the proportion of hits from Chromalveolates species for each of the 2 473 domains (see Figure 5.(b)). We found that 326 domains had a strong majority (> 90%) of hits from Chromalveolates species and hence could be considered as specific to this super-group.

**Figure 5.**
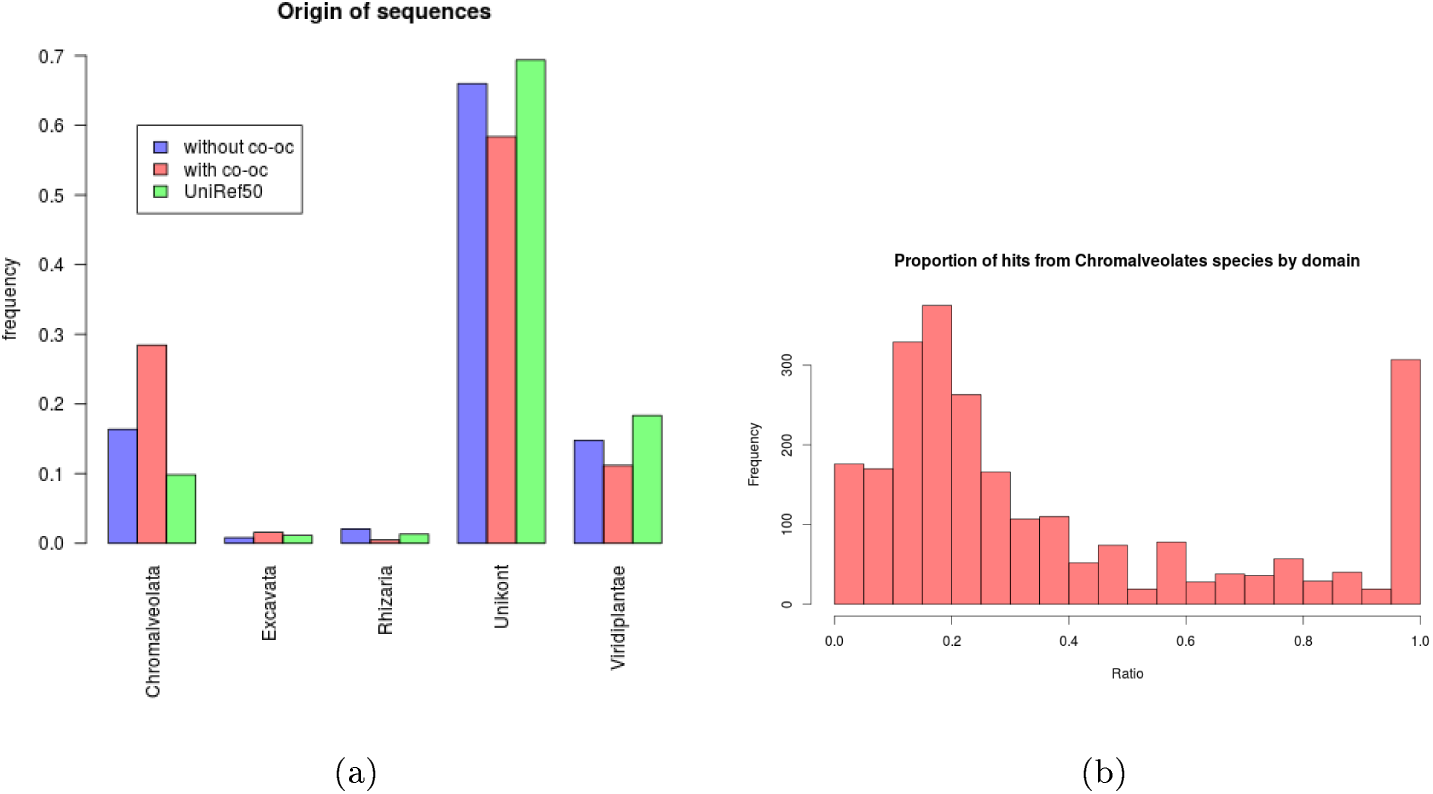
(a) Hit distribution among the five eukaryotic super-groups (Keeling et al, 2005). In green, the distribution in UniRef50 (restricted to eukaryotic sequences); in blue the distribution of all BLAST hits; in red the distribution of hits selected by cooccurrence. (b) Distribution of the proportion of Chromalveolate hits in the new families.

#### Domain assessment

We next investigated the quality of the newly discovered domains. In all the following, we use the domains identified with the procedure integrating the already known Pfam domains. We used different quality measures (see below) and compared our results to the same measures applied to Pfam domain families. In order to compare families built on similar sequences, we restricted our analysis to Pfam families present on a *P. falciparum* protein, and the associated MSAs were restricted to sequences from Uniref50. Moreover, to avoid any bias due to the alignment step, Pfam families were realigned using the same tool and parameters we used to produce our models. Next, in order to assess the benefits of using co-occurrence information, we also conducted the same experiments on domains that were predicted using the total hit density instead of the co-occurring hit density. Namely, we used the same iterative procedure for identifying hit clusters, but applied it directly to the hit density (in blue on Figure 3).

It is hard to assess the quality of a multiple sequence alignment in the absence of other information. We propose to use four types of measures for this. The first measure (Figure 6(a)) is the alignment *homogeneity.* A good alignment usually has a minority of insertions and deletions on each position. This can be measured by the proportion of residues on each position. Namely, this proportion should be either very low (when a few sequences have an insert at this position) or high (when a few sequences have a deletion at this position). We measure the homogeneity by

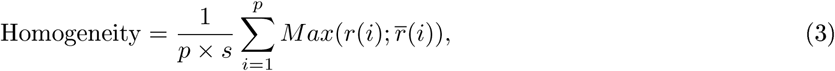

with *s* and *p* representing the number of sequence and the length of the MSA, respectively, while *r*(*i*) and *r̅*(*i*) represent the number of residues and *indels* (insertion or deletion) at position *i*, respectively. With this formula, a good alignment should have a score that tends to 1.

**Figure 6.**
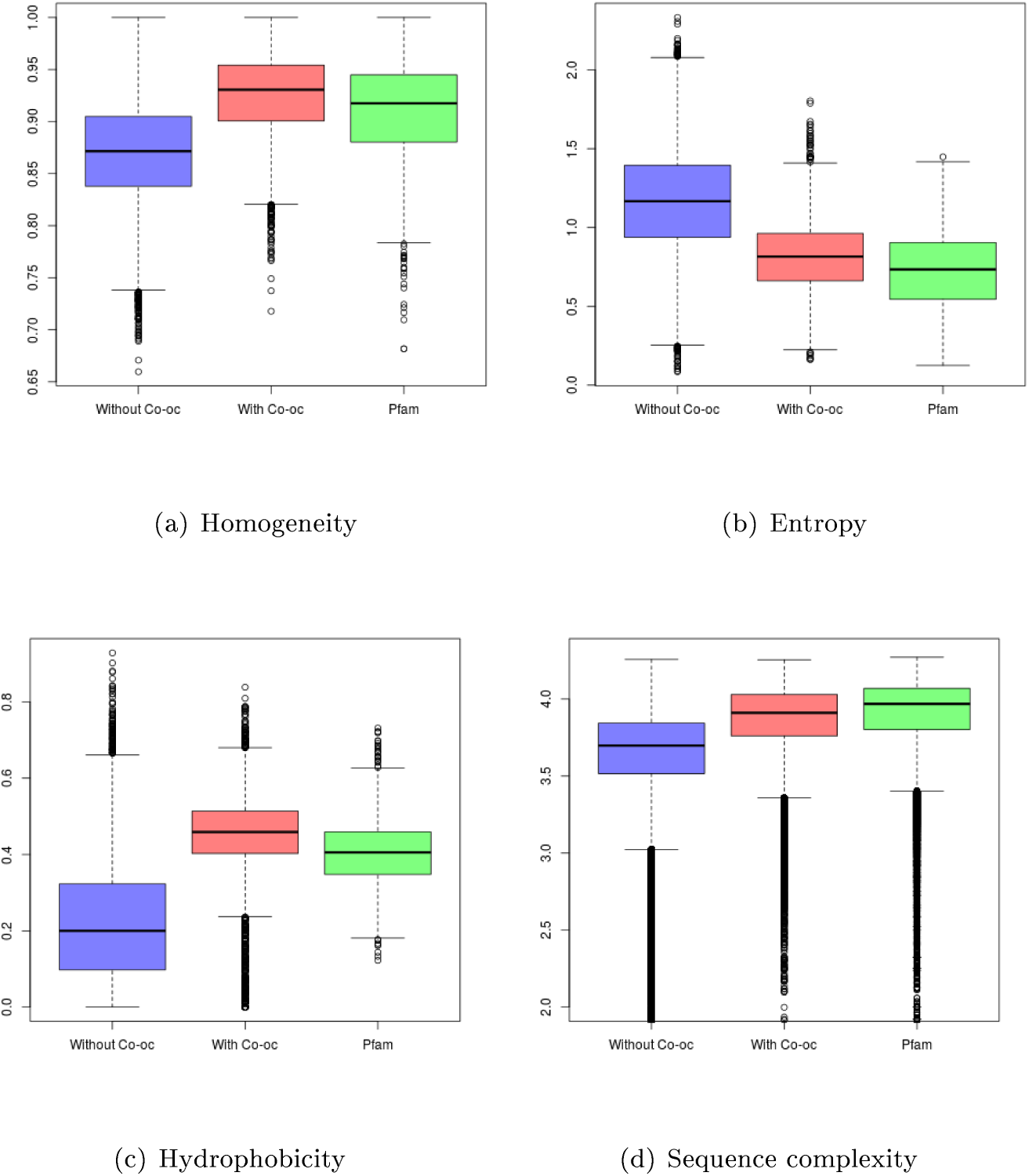
Quality scores measured on models obtained without co-occurrence (in blue), models obtained with co-occurrence (in red) and Pfam models (in green).

The second measure (Figure 6(b)) is the *entropy* of match states in the alignment. Entropy is based on amino-acid classes. Briefly, amino acids can be grouped according to physicochemical features. We used the same definition of classes as in the Seaview software package (Gouy et al, 2010).

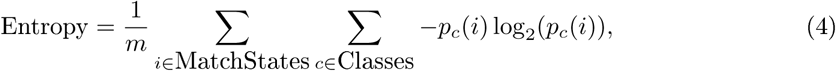

with *m* representing the number of match states in the MSA. We consider a position as a match state if the number of residues at the position is higher than the number of indels. *p_c_*(*i*) represents the proportion of residues belonging to class *c* at position *i*. In a good alignment, all residues on a match state tend to belong to the same amino-acid class and the entropy tends to 0. In a poor alignment, the distribution of residue classes is equi-probable and the entropy tends to ≈ 3.

The third measure (Figure 6(c)) is the *hydrophobicity* score:

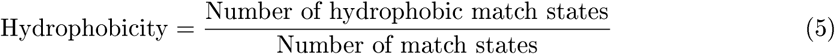

We consider a match state as hydrophobic if the majority of residues at this position are considered as hydrophobic (residues L, A, F, W, V, Μ, I, P, C and G). The hydrophobic residue proportion is commonly used as a measure of globularity, because globular domains have a stable amount of strong hydrophobic amino acids (about one third of the sequence) (Dill, 1985). Note that contrary to homogeneity and entropy it is difficult to establish what would be a “good” hydrophobic score. So here we will essentially compare our results to that of Pfam domains.

The last measure (Figure 6(d)) is the *complexity* of sub-sequences in the MSA.

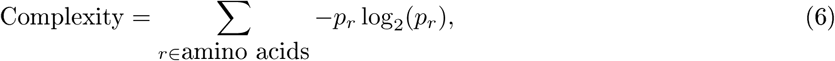

with *p_r_* representing the relative frequency of amino acid *r* in the sequence at hand. Contrary to the three previous measures, which are based on the columns (positions) of the alignments, this one is applied on each sequence independently. Measuring the sequence complexity is a useful way of identifying repetitive sequences which are characteristic of non-globular regions (Wootton, 1994).

As we can see on Figure 6, domain families identified with co-occurrence obtain quality scores relatively close to that of standard Pfam families. On the contrary, results obtained with all BLAST hits without the co-occurrence filtering are far from the standard Pfam families. This illustrates the interest of the approach and show that co-occurrence provides a useful way to filter out results of BLAST search at genome scale.

#### Domain comparisons

We next assessed if some of the new domains were similar to known Pfam domain families that would not have been identified on the *P. falciparum* proteins (remember that the known Pfam occurrences have been masked in this experiment), or to other newly discovered domains (the same family can be identified several time on given proteome). We thus ran several profile/profile comparisons using HHPred (Soding et al, 2005). As above, we computed, for each HMM alignment, a p-value and an overlap ratio between the two HMMs. We first ran an all versus all comparison of Pfam HMMs. We identified, for each Pfam HMM, the other Pfam HMM that most resembles it. Figure 7(a) reports the p-values and overlap ratios of this analysis. As we can see, most Pfam HMM pairs have a p-value > 10^−10^ and/or an overlap ratio below 0.8. Hence we used these two cutoffs as a rough criterion to decide whether two HMMs are similar or not. Figure 7(b) reports the results of the comparisons between our new HMMs and Pfam HMMs. With the above criteria, we found that, on the 2 473 models we produced, only 168 are strictly similar to a Pfam model and hence likely constitute an undetected occurrence of a known Pfam domain family. We noted however that a large part of our models get p-values below 10^−10^, indicating that they locally resemble a Pfam family. While such local resemblances is quite common between Pfam families themselves (see the numerous points in the top-left quarter of Figure 7(a)) it is possible that part of these new domains are actually partial occurrences of already known Pfam families. Genuine partial occurrences seems quite rare events (Prakash and Bateman, 2015), but it is known that alignment and annotation artifacts may result in partial domain observation (Triant and Pearson, 2015). To test this hypothesis, we compared the size of the new domain to that of the Pfam family that most resembles it. Surprisingly for the 1955 domains which have good p-value (< 10^−10^) but low overlap (< 80%) with a Pfam family, the new domain is longer than the Pfam family in 89% of the time. Moreover, most of the time the Pfam model is very short (see Figure 7(c)). Interestingly we observe the same type of short domains in the Pfam vs. Pfam comparison when restricting to domain pairs with good p-value and low overlap (Figure 7(c)). Hence, rather than partial domains, the newly identified domain could rather be viewed as extensions of smaller Pfam domains, something already quite common in the Pfam database. Finally, we compared our models with each other (Figure 7(d)). 1 215 of the new models did not seem to have close similarities with other new models, while 1 258 models were similar to one or more other new models. This higher proportion of redundancy compared to that observed against the Pfam database was somewhat expected, because domain families often have multiple occurrences in one proteome (Vogel et al, 2005).

**Figure 7.**
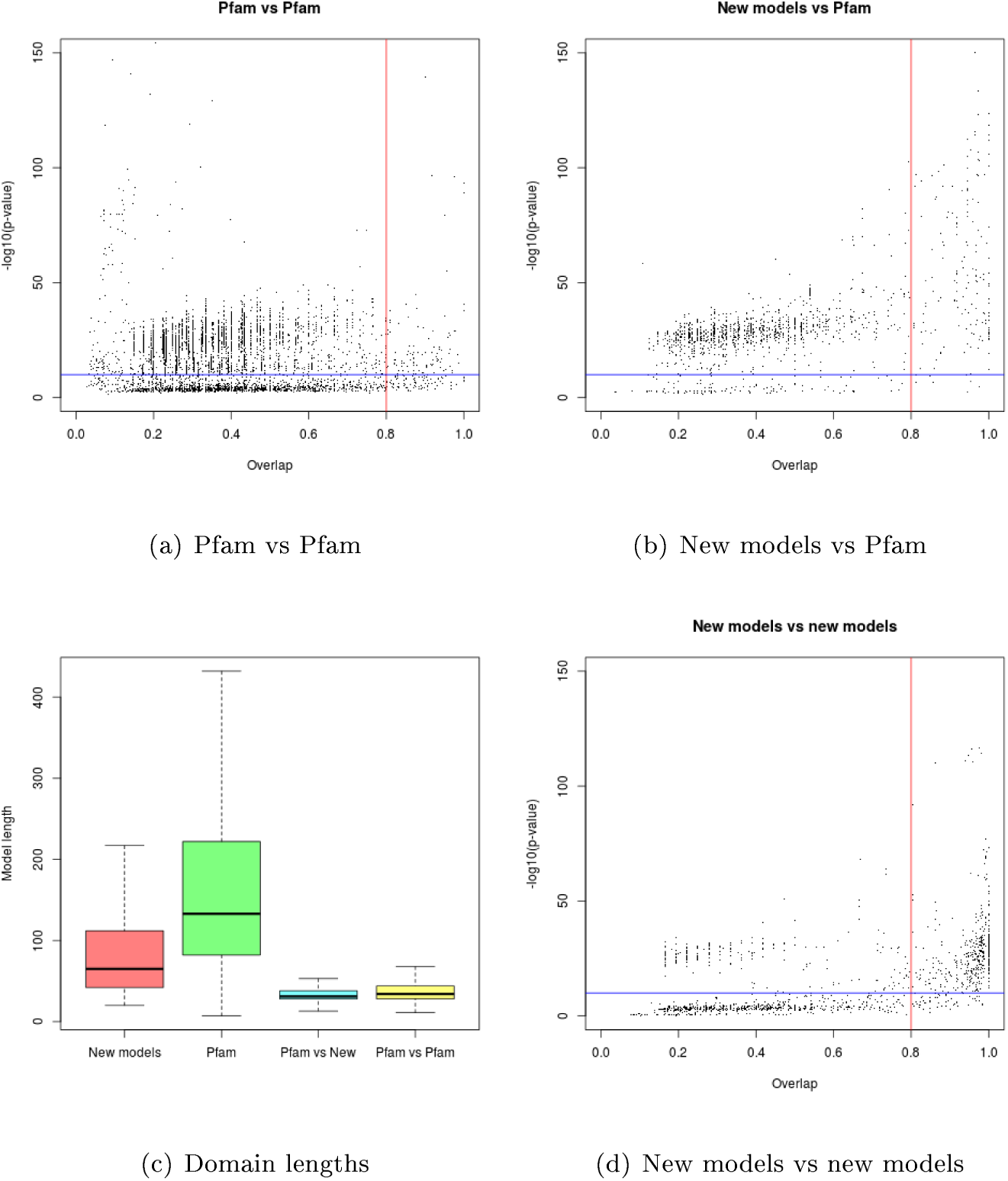
HMM/HMM comparison of new domain families and Pfam domain families. In Figures (a), (b) and (d), each point is associated with one particular HMM and corresponds to the best alignment found between this and all other HMMs. The x-axis shows the overlap ratio of the local alignment between the two HMMs; the y-axis indicates the negative log. of the alignment p-value; blue line corresponds to *y =* –*log*(10^−10^); while the red line corresponds to *x =* 0.8. Figure (c) shows the lengths of the models obtained by our approach (red), the lengths of all Pfam models (green), the lengths of the Pfam models associated with the points depicted in the top left quarter of figure (b) (blue), the length of the smallest Pfam model associated with the points depicted in the top left quarter of figure (a) (yellow)

#### Comparison against other automatically generated databases

We compared the results obtained by our procedure to Pfam-B and ProDom (Servant, 2002), two automatically generated domain databases. Pfam-B was part of the Pfam database until release 26. Actually, Pfam contained two types of domain families until this release: the high quality and manually curated Pfam-A families (“classical” Pfam domains, which are usually and until here in this article just called “Pfam”), and Pfam-B families, which were automatically generated by the ADDA algorithm (Heger and Holm, 2003) on the basis of all parts of Uniprot sequences not already covered by a Pfam-A occurrence. ProDom is a protein domain family database constructed automatically by clustering homologous segments with the MKDOM2 procedure based on recursive PSI-BLAST searches on the Uniprot database. Among the 460125 families contained in Pfam-B (release 26), we found 651 families with at least 5 different proteins of the UniRef50 database. These 651 families cover about 9.70% of the *Plasmodium falciparum* proteome. The ProDom database (December 2015 release) contains numerous very small families less than 30 nucleotides long. To be consistent with our experiments and the Pfam database, these families were not considered in the following. Among all 3 7391 57 families contained in ProDom, we found 1453 families longer than 30 nucleotides, with at least 5 proteins of the UniRef50 database, one protein of *P. falciparum,* and that did not overlap with a Pfam-A domain. These families cover about 3.99% of the *P. falciparum* proteome. For comparison, clusters with at least 5 hits identified by our approach cover 7.12% of *P. falciparum* residues. Hence, in terms of coverage, our approach (7.12% lies between ProDom (3.99% and Pfam-B (9.70%).

We realigned these families using the same tool and parameters used to produce our models, and for each approach we computed the quality scores presented above (see Figure 8). As we can see, ProDom, and more importantly Pfam-B families, have very low homogeneity scores compared to Pfam-A families. This illustrates the fact that these databases sometimes include in one family different sequences that cannot be aligned and that should be clustered apart. On the contrary, they have good entropy scores, even better than that of Pfam-A families. This, however, is a mechanistic effect of the very low number of sequences that compose these families when they are restricted to UniRef50 sequences, as illustrated in Figure 8 (e). For Pfam-B, for example, 80% of the considered families have less than 15 sequences in UniRef50. Hence, it seems that many families include only highly similar sequences, which obviously provides good entropy scores, but also induces a lack of diversity. On the contrary, by using co-occurrence information, our approach allows selection of more diverse sequences without the issue of introducing more false positives. In terms of hydrophobicity, ProDom has scores comparable to that of our approach and Pfam-A, while Pfam-B achieves slightly lower scores. Finally, ProDom and Pfam-B have comparable sequence complexity, but slightly lower than that of our approach and Pfam-A. Altogether, these results suggest that the domains built using co-occurrence are globally of better quality than those found in Pfam-B and ProDom. Note, however, that the purpose of these two databases is different from and somewhat wider than ours. Indeed, their goal is to build a library that pools all domain families of every species, while our aim is to find new domain occurrences for a specific species (here *P. falciparum*).

**Figure 8.**
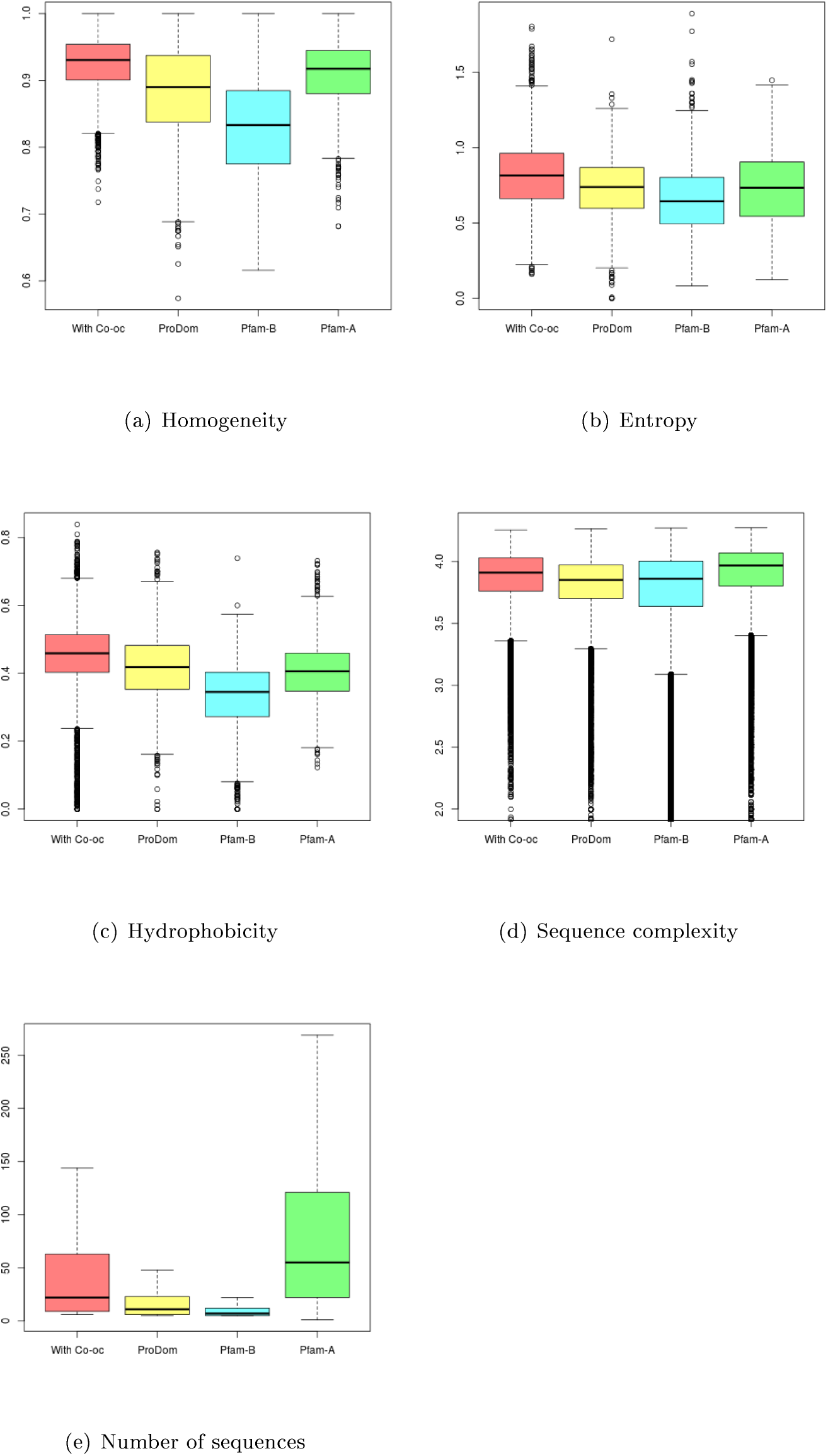
Quality scores (a-d) measured on families obtained by our approach (in red), ProDom families (in yellow), Pfam-B families (in cyan) and Pfam-A families (in green). Figure (e) shows the number of sequences in the families of the different databases.

#### Test against previous release

As an ultimate test, we ran our analysis using an older version of Pfam (release 26) and computed the number of new domains of the 28 release that this 26 analysis allowed us to recover. Over the 92 domains referenced in Pfam 28 and not in Pfam 26, we identified a domain on the same region for 54 of them. Careful inspection of these domains revealed that the new domains often have strong similarity with the domains identified by our approach. For example, the domain we identified on protein 077317_PLAF7 shows high similarity with the domain family PF16876 of the 28 release (p-value = 4 * 1*e*^−31^ and overlap = 0.99).

#### Gene ontology annotations

The Gene Ontology (GO) Consortium (Ashburner et al, 2000) provides a structured vocabulary describing gene functions according to three points of view (biological process, molecular function, and cellular component). Each ontology is organized as a directed acyclic graph where each node is associated with a term, while edges describe specialization and generalization relationships. The GO Consortium also provides a list of protein annotations for most sequenced organisms. With this information, we tried to associate an annotation with the new domains identified by our procedure. For each domain (cluster), we gathered annotations associated with proteins from where the different hits belong to (cluster members). Then we parsed the GO, and if more than 95% of annotated sequences shared the same GO term, we annotated the domain with this function. We set a minimum of 5 annotated sequences in a cluster to avoid irrelevant annotations. GO terms that are direct children of the root node were not considered as significant annotations and were ignored. Among the 2 473 newly identified domains, we can thus propose an annotation for 1 394 domains (see Supp. Data). Among these, for 1 273 domains the proposed annotation is consistent with already known annotations of the *P. falciparum* protein where the domain has been identified. For 121 domains, the proposed annotations extend the known annotations.

To go one step further, we tried to identify other occurrences of our new families in the *P. falciparum* proteome. Because of duplication events, it is quite usual to find multiple occurrences of the same domain in a proteome (Vogel et al, 2005). We ran a HMMScan of our 2 473 new families against *P. falciparum* proteins with an e**-**value threshold of 10^−10^. If two hits overlapped, only hits with the best e-value was kept. This way, we were able to identify 843 new domain occurrences on 671 different proteins (with 638 different models) that were not covered by a Pfam or one of our previously identified domains. These new occurrences allowed us to propose new annotations for 34 additional proteins.

## Discussion and conclusion

Here we proposed a new method to take co-occurrence into account in a typical BLAST analysis and to construct new domain families on these results. Our method is based on the analysis of co-occurring hit density along the query protein. We designed a clustering procedure to identify clusters of similar hits that sign domain boundaries and a statistical test to assess the relevance of the identified clusters. Moreover, we have presented a procedure to estimate the proportion of false positives in a set of clusters.

We used *Plasmodium falciparum* as a case study to evaluate our approach. Our experiments showed an increase of 16% of the number of significant hits and an increase of 28% of the covered proteome. We identified 8 530 significant hit clusters. The false detection rate was estimated at around 5%. We used these clusters to construct new domain families and to enrich databases of known domains. These models showed quality close to that of standard Pfam models and quite moderate redundancies with respect to original Pfam models, which indicates that they likely belong to new domain families that are not yet referenced in the database. Conversely, we identified more redundancies among the generated models themselves, which indicated the presence of multiple occurrences of a same domain family in the *P. falciparum* proteome.

Our approach could be improved in several ways. First, clustering is done by an ad-hoc procedure which involves two parameters that are set empirically. Hence an interesting improvement would be to integrate an automatic procedure able to find parameter values that best fit the protein at hand. Similarly, the quality of generated HMMs depends on the parameters of the alignment softwares (here BLAST and MUSCLE). We used standard parameter values in this study, but other parameters could certainly help in building better models. This is especially true for species like *P. falciparum,* which have amino-acid distributions far from the classical distribution observed in other species. Finally, the main drawback of our approach is that it cannot annotate all proteins of a given species. For *P. falciparum,* for example, our procedure is unable to identify any cluster in a total of 1 685 proteins. The main reason for this is the existence of mono-domain proteins for which domain co-occurrence is of no use. In this case, it would be interesting to identify other features that could replace domains in the co-occurrence analysis.

For example, tandem repeat sequences or disordered regions that constitute other classes of conserved protein sequences may be helpful in certain cases.

## Methods

### Identifying domains from BLAST hits

We used BLASTP software (Altschul et al, 1990), release 2.2.28 with default parameters and a max e-value set at 10^−2^. All hits smaller than 30 residues were removed, and in case of overlapping hits on a target protein only the hit with the lowest e-value was considered. Then hits were clustered according to algorithm 1. Each cluster corresponded to a putative domain.

~~~
**Algorithm 1** Clustering of similar hits
**Input:** *ℋ*: a set of co-occurrent hits on a query protein
**Output:** *𝒞* a set of hit clusters. Each cluster *C* ∈ *𝒞* is defined by a set of hits and a start *C_s_* and
    end *C_e_* position on the protein.
    *𝒞* ← 0
    **do**
           *ℋ*\* — *ℋ* minus all hits overlapping any cluster *C* ∈ *𝒞*
       Compute hits density with *ℋ*\*
       *P* ← position with highest density
       *C* ← all hits covering P
       *C*_s_ ← lowest starting position in *C*
       *C_e_* ← highest ending posi ion in *C*
       **do**
            Unstable ← False
            *N* ← *C* size
            Compute *Ns* and *Ne*, the number of hits in *C* covering *C_s_* and *C_e_*, respectively
            **if** *Ns* **<** 1/3 · *N* **then**
                *C_s_* ← *C_s_* + 1
               Unstable ← True
            **end if**
            **if** Ne **<**1/3 *N* **then**
                *C_e_* ← *C_e_* – 1
                Unstable ← True
            **end if**
            Remove hits in *C* if more than 30% of residues outside range [*C_s_* … *C_e_*]
        **while** Unstable
        **if** (*P_e_* — *P_s_***) >** 30 **then**
               Add *C* in *𝒞*
         **end if**
    **while** at least one new cluster in *𝒞*
~~~

### FDR estimation

First, our approach is run on all proteins of the query organism, and the number of domains below a given p-value threshold is computed. Then, all BLAST hits are randomly shuffled among all proteins. To preserve the homology relationship between hits during the randomization process, the hits are not shuffled independently among the proteins but rather by entire clusters. We thus use the clustering computed from the co-occurring hit density. All hits with no co-occurrence are added to the cluster they overlap if less than 20% of their residues are outside the cluster. These clusters of hits are then randomly permuted among proteins. This creates a situation where each cluster loses its previous co-occurrences but may fortuitously find new ones on the new protein. Once hits have been randomly distributed on proteins, the number of co-occurring hits are computed in each cluster, and a p-value is estimated with Formula (1). The number of clusters below the chosen p-value threshold is computed, and the entire procedure is resumed several times (*e.g.* 10 times) to get a better estimate of the number of domains that can be identified in random data. This number is then used to estimate the FDR of the procedure with Formula (2).

### HMM learning

HMM are trained using HMMER3 software (release 3. 1b2) with the following command:

hmmbuild -n <hmm__name> amino fast <hmmfile_out> <msafile> ; with <hmm_name> a name for the trained HMM, amino specifies that input alignement is protein sequence data, fast assigns columns with >= 50% (default) residues as match position, <hmmfile_out> the output HMM file, <msafile> the input alignement.

### HMM/HMM comparisons

HMM-HMM comparisons are done using the hhsearch software provided in the hhsuite (release 2.0.16) with the following command:

hhsearch -i <hmmfile_in> -d <hmm_db> -o <resfile_out> -loc with <hmmfile_in> the input HMM, <hmm_db> the HMM database to compare to, <resfile_out> the file containing results, -loc to search the best local alignement.

## Author's contributions

Text for this section …

## Acknowledgements

We thank Cédric Notredame for helpful discussions. We thank the ProDom team for providing us with the last version of the ProDom database.

## Additional Files

### Additional file 1 — New domain families identified in *P. falciparum*

Pfalciparum_new_families_fasta.zip

This zip file contains the new domain families identified in *P. falciparum.* Each family is a multiple alignments (fasta format) of the BLAST hits (UniRef50) that have been used to construct the family.

### Additional file 2 & 3 — HMMs of new domain families identified in *P. falciparum*

Pfalciparum_new_families_HMM.hmm & Pfalciparum_new_families_HMM.hhm Both files contain the HMMs (either in HMMER or HHPRED format) built from the families containing at least 5 sequences.

### Additional file 4 — GO annotations of new domain families identified in *P. falciparum*

Pfalciparum_new_families_go_annotations.tsv

This file contains the GO annotations that have been identified for the families containing at least 5 sequences.

